# Balanced excitation and inhibition decorrelates visual feature representation in the mammalian inner retina

**DOI:** 10.1101/040642

**Authors:** Katrin Franke, Philipp Berens, Timm Schubert, Matthias Bethge, Thomas Euler, Tom Baden

## Abstract

The retina extracts visual features for transmission to the brain. Different types of bipolar cell split the photoreceptor input into parallel channels and provide the excitatory drive for downstream visual circuits. Anatomically, mouse bipolar cell types have been described down to the ultrastructural level, but a similarly deep understanding of their functional diversity is lacking. By imaging light-driven glutamate release from more than 11,000 bipolar cell axon terminals in the intact retina, we here show that bipolar cell functional diversity is generated by the balanced interplay of dendritic excitatory inputs and axonal inhibitory inputs. The resultant centre and surround components of bipolar cell receptive fields interact to decorrelate bipolar cell output in the spatial and temporal domain. Our findings highlight the importance of inhibitory circuits in generating functionally diverse excitatory pathways and suggest that decorrelation of parallel visual pathways begins already at the second synapse of the mouse visual system.

## INTRODUCTION

The retina is the first processing stage of the visual system. It extracts features like motion or edges (Wässle, 2004; Masland, 2012a) and relays these to the brain through a highly diverse set of retinal ganglion cells (RGCs) (Sanes and Masland, 2015; Baden et al., 2016). This functional diversity starts to emerge already at the first retinal synapse, where the visual signal is distributed from the photoreceptors onto ∼14 bipolar cell (BC) types (reviewed in (Euler et al., 2014)). Their axon terminals stratify at different depths of the inner plexiform layer (IPL) and provide the excitatory drive for the feature extracting circuits of the retina.

Anatomically, the set of mouse BC types is well characterised (Ghosh et al., 2004; Wässle et al., 2009b) and their exact number and ultrastructural connectivity are known (Helmstaedter et al., 2013; Kim et al., 2014). Functionally, BCs have been classified mostly into broad categories like On and Off, transient and sustained or chromatic and achromatic (Euler et al., 1996; Awatramani and Slaughter, 2000; DeVries, 2000; Li and DeVries, 2006; Breuninger et al., 2011); however, a deeper understanding of their functional diversity and its origin is lacking.

Some of the observed differences between BC types including polarity and chromatic selectivity are established in the outer retina through differences in the excitatory dendritic input from photoreceptors (DeVries et al., 2006; Puller et al., 2013; Borghuis et al., 2014; Lindstrom et al., 2014; Puthussery et al., 2014). In the inner retina, more than 40 types of inhibitory amacrine cell (AC) modulate BC output at the level of their synaptic terminals (Strettoi et al., 1990; Masland, 2012b; Helmstaedter et al., 2013). While a handful of AC circuits have been studied at great detail (e.g. the A17 (Grimes et al., 2009) or the AII network (Demb and Singer, 2012)), we do not understand the general principles by which AC circuits help to decompose the visual scene into the parallel channels carried by the BCs. This fundamentally requires recording from many BC terminals in the retina with long-range connections preserved.

To address this important aspect, we took advantage of the recently developed glutamate biosensor iGluSnFR (Borghuis et al., 2013; Marvin et al., 2013) and set out to systematically characterise the complete glutamatergic output of mouse BCs at the level of individual axon terminals in the whole-mounted retina. In contrast to presynaptic calcium, which has been used to assess BC function in mouse and zebrafish (Dreosti et al., 2009; Baden et al., 2012; Yonehara et al., 2013; Chen et al., 2014a), glutamate release represents the output “currency” of BCs, not only accounting for presynaptic inhibition but also any release dynamics of BC ribbon synapses (Burrone and Lagnado, 2000; Cho and von Gersdorff, 2012; Nikolaev et al., 2013). Combining systematic functional population recordings with available anatomical information provides a rare opportunity to understand an entire class of neurons at unprecedented depth.

## RESULTS

### Glutamate release units of the IPL

To survey light-driven glutamate release across the IPL of the whole-mounted mouse retina we used time-lapsed two-photon imaging of the fluorescent glutamate biosensor iGluSnFR (Borghuis et al., 2013; Marvin et al., 2013). Intravitreal injection of AAV9.iGluSnFR directed at the naso-ventral retina (Fig. 1a, Methods) yielded homogenous ubiquitous expression throughout the IPL (Fig. 1b), thus allowing for sampling of glutamate release at all IPL depths (SFig. 1f). For each scan field, we registered the recording depth as its relative distance to the two plexi of SR101-stained blood vessels. The error in estimating IPL depth was <1 μm, as verified by comparison to TdTomato-positive ChAT bands (Famiglietti and Tumosa, 1987) (Fig. 1c, Methods).

**Figure 1.**
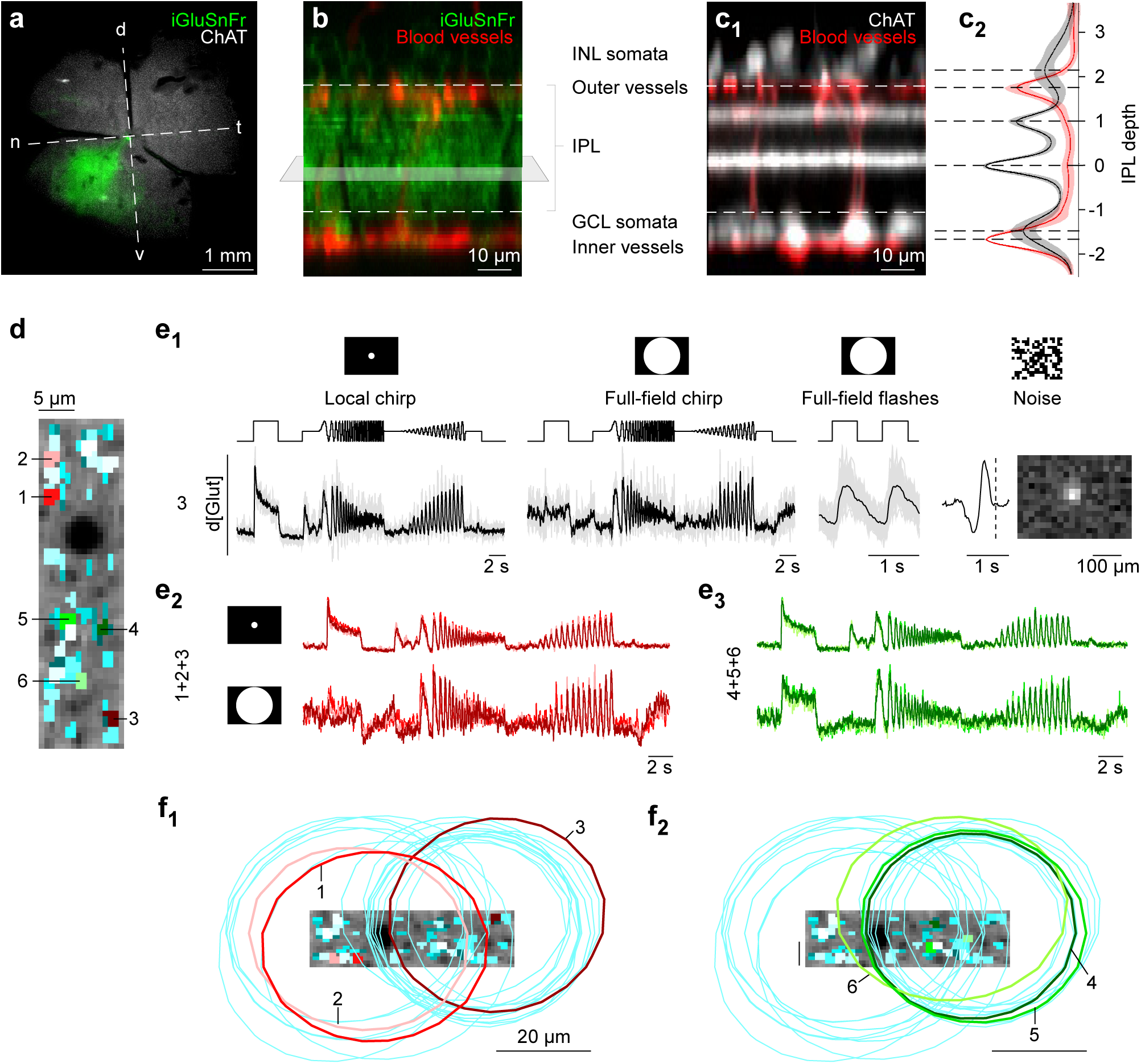
Imaging light-driven glutamate release in the IPL. **a**, Whole-mounted retina of a ChATCre x AI9^tdTomato^ mouse (tdTomato in *white)* with ubiquitous iGluSnFR expression (*green*) in the naso-ventral retina following intravitreal injection of <u>AAV9.hSyn.iGluSnFr.WPRE.SV40</u>. d: dorsal, v: ventral, n: nasal, t: temporal. Unless indicated otherwise, vertical and horizontal scales are identical. **b**, Vertical projection of a high resolution stack showing iGluSnFR expression (*green*) across the IPL, with blood vessels visualised by SR101 (*red*). Grey plane illustrates scan field orientation. GCL: Ganglion cell layer, IPL: Inner plexiform layer, INL: Inner nuclear layer. **c**, Position of ChAT bands (*white*) relative to blood vessel plexi (*red*, c_1_) and their average depth profiles (± s.d. shading, c_2_); n=9 stacks from 3 mice. **d**, Example scan field (64×16 pixels) with Region-of-interest (ROI) mask overlaid. **e**, Glutamate traces of one ROI from (d) in response to (from left to right) local and full-field chirp stimulus, full-field flashes, temporal and spatial receptive field (e1, Methods). Glutamate traces represent relative glutamate release (d[Glut]). Individual trials in grey (n=5 for chirps, n>30 for full-field flashes) with mean responses overlaid (*black*). Dotted line in temporal RF indicates time point of response. e_2_,e_3_, Superimposed mean glutamate traces in response to local (*top*) and full-field chirp (*bottom*) of red (e_2_) and green (e_3_) ROIs from (d). **f**, Scan field and ROI mask from (d) with spatial RFs (2 s.d. outlines of gauss fit, n=20 ROIs) of red (f1) and green (f_2_) ROIs color-coded in (d).

To objectively define individual glutamate “release units”, we placed regions-of-interest (ROIs) in a single scan field (typically 48x12 μm at 32.125 Hz) using a custom algorithm based on local image correlation over time (Fig. 1d, SFig. 1, Methods). This allowed us to sample light-driven activity of 74±24 ROIs per scan field (SFig. 1e, SVid. 1). We verified the performance of this algorithm using calcium imaging of BCs with the GCaMP6f biosensor (Chen et al., 2013), where individual axon terminals could be easily resolved (SFig. 2a). Our algorithm reliably detected individual terminals and rather assigned two ROIs to a single terminal before merging two terminals into one ROI (SFig. 1d, SFig. 2a). In addition, receptive field (RF, see below) sizes estimated from calcium signals of single terminals closely fit those estimated from single iGluSnFR ROIs (SFig. 2c,d) and matched the anatomical dimensions of BC dendritic fields (Wässle et al., 2009b; Behrens, Schubert, Haverkamp, Euler and Berens, personal communication). Accordingly, each ROI likely captured the light-driven glutamate signal of at most one BC axon terminal.

We used a standardised set of four light stimuli (Fig. 1e_1_, see also (Baden et al., 2016)) to characterise BC output across the IPL: (i) local (100 μm diameter) and (ii) full-field (600x800 μm) “chirp”-stimuli to probe response polarity as well as contrast and frequency preference of BC centre and centre-surround, respectively, (iii) 1 Hz full-field steps to study response kinetics and (iv) binary dense noise to estimate receptive fields (Methods).

ROIs in a single scan field could typically be grouped into two or more distinct response profiles (e.g. Fig. 1e_2,3_, red versus green ROIs), suggesting that multiple BC types could be recorded at a single IPL depth as expected from their partial stratification overlap (Greene et al., n.d.; Helmstaedter et al., 2013; Kim et al., 2014) (cf. Fig. 2a). Similarly, the reported tiling of the retinal surface by neurons of the same type implies that more than one cell of a BC type may contribute terminals to a single scan field (Wässle et al., 2009b). Indeed, ROIs that shared a common response profile had RFs that either almost completely overlapped or were spatially offset consistent with BC tiling (Wässle et al., 2009b) (Fig. 1f). For example, the highly overlapping spatial RFs of the green ROIs suggest that they correspond to terminals not only belonging to the same type of BC, but the same cell (Fig. 1f_2_, see also SFig. 2a-c). In contrast, the red ROIs likely correspond to terminals of two neighbouring cells of a second type (Fig. 1f_1_). Taken together, we therefore think that our ROIs reflect a reliable measure of BC output at the level of individual axon terminals, the computational output unit of the inner retina.

**Figure 2.**
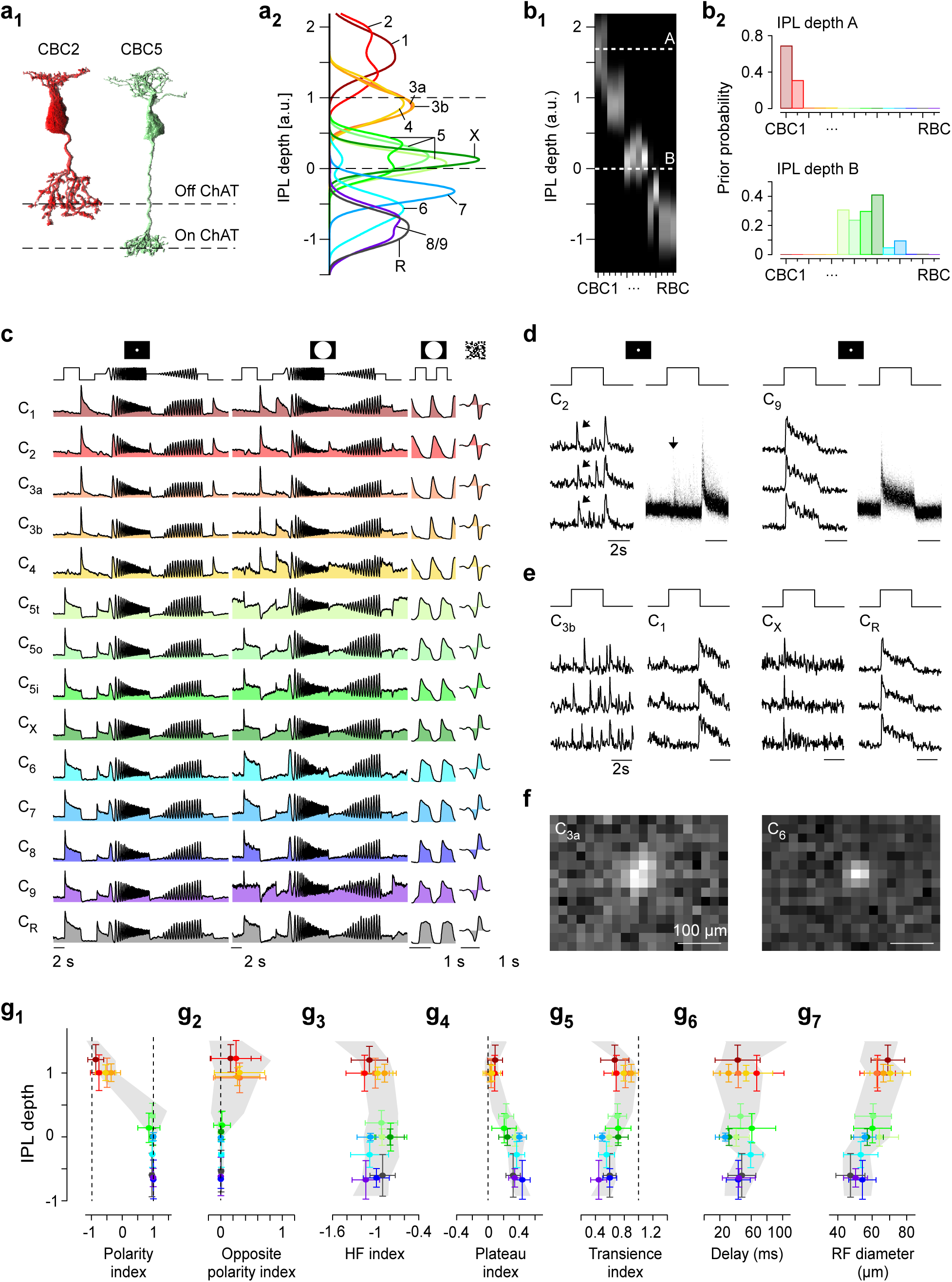
Anatomy-guided clustering and functional organisation of the IPL. **a**, EM-reconstructed and volume-rendered example BCs (**a1**, from (Helmstaedter et al., 2013)), illustrate type-specific axonal stratification patterns relative to ChAT bands (*dashed lines*). a_2_, Mean BC stratification profiles of all known BCs (5 Off CBCs, 8 On CBCs, 1 rod BC) (Greene et al., n.d.; Kim et al., 2014). **b**, Exemplary distributions of prior probabilities for cluster allocation (b_2_) taken from mean stratification profiles (b1) of scan fields recorded at two different IPL depths (A: 1.7; B: 0). **c**, Normalised mean glutamate response of every cluster to (from left to right) local and full-field chirp, full-field flashes and temporal kernels estimated from the noise stimulus. **d**, Glutamate responses to the local chirp step stimulus of single ROIs assigned to C_2_ and C_9_, respectively. Shown are responses to 3 trials and a histogram of response amplitudes across each cluster’s 100 best responding ROIs. On responses in Off BC cluster C_2_ are highlighted (*arrows*). **e**, As (d), showing “spiking” (C_3b_, C_X_) and non-spiking (C_1_, C_R_) responses in clusters of either polarity. **f**, Spatial RFs of individual ROIs assigned to C_3a_ and C_6_, respectively. **g**, Response measures estimated for all ROIs (n=8,448) plotted against IPL depth, with grey shading corresponding to median ± s.d. for every IPL bin (n=13 bins). Cluster means (± s.d.) are overlaid. From left to right: Response polarity (g1), opposite polarity events (g_2_), HF index (g_3_), plateau index (g_4_), response transience (g_5_), response delay (g_6_) and RF diameter (g_7_). g_1-6_ were estimated from local chirp step responses and g_7_ was estimated from spatial RFs obtained from the noise stimulus.

### Anatomy-guided functional clustering of mouse BCs

In total, we recorded light-evoked BC glutamate release from 11,101 ROIs (n=150 scan fields, n=29 mice) throughout the IPL, each scan field tagged to its precise IPL depth. For the following analysis, we assumed that (i) BCs are the main source of glutamate in the inner retina (Discussion) and (ii) the anatomical catalogue of BCs in the mouse retina is complete at a count of 14 types (5 Off cone BCs (CBCs), 8 On CBCs and rod BC (RBC)), each forming a complete and independent mosaic (Wässle et al., 2009b; Helmstaedter et al., 2013; Kim et al., 2014; Greene et al., 2016). Following this assumption, the measured glutamate signals must map onto these 14 types of BCs including RBCs (SFig. 2e,f). We took advantage of available EM-reconstruction data on BC axonal stratification profiles (Kim et al., 2014; Greene et al., 2016) to guide a functional clustering algorithm (Fig. 2a): For each scan field taken at a specific IPL depth, a prior probability for cluster allocation was computed from the relative axon terminal volume of all BC types in the respective IPL stratum (Fig. 2b). For example, all ROIs of a scan field taken at an IPL depth of 1.7 were likely to be sorted into clusters for CBC types 1 and 2 (Fig. 2b_2_, top), while a scan field taken at a depth of 0 received a bias for CBC types 5-7 (Fig. 2b_2_, bottom). We then extracted functional features from the glutamate responses that passed our quality criterion (76.1% of ROIs) using sparse PCA (SFig. 3a) and clustered the ROIs using a modified Mixture of Gaussian model (Methods).

This yielded a functional fingerprint of every anatomical BC type in the mouse retina (Fig. 2c, SFig. 3b), including a detailed account of their response kinetics, RF structure and centre-surround properties. ROIs allocated to each cluster originated from at least 10 scan fields and 5 animals. Functional clusters were well-separated in feature space including pairs of the same polarity (SFig. 3c) and individual scan fields routinely comprised ROIs allocated to more than one cluster (SFig. 3d). Because some BC types have highly overlapping stratification profiles, the assignment of our functional cluster to morphological types is not bijective. This caveat pertains to clusters (C): C_1,2_, C_3a,b,4_, C_5a-c,X_ and C_8,9,R_ (SFig. 3b), which might need to be permuted. For simplicity, we refer to functional clusters by the anatomical BC profiles they originated from.

### Organisational principles of the IPL

A fundamental principle of vertebrate inner retinal organisation is the subdivision into Off and On cells (Werblin and Dowling, 1969; Nelson and Kolb, 1983; Euler et al., 1996). In agreement, C_1-4_ increased activity at the offset of a step of light (Off BCs), while C_5a-R_ responded to the step’s onset (On BCs, Fig. 2g_1_). However, the segregation into On and Off responses was not as clear-cut as expected: All Off BC clusters frequently responded with delayed spike-like events during the On phase of the light step (Fig. 2d,g_2_). These On events in Off BC clusters did not correspond to spontaneous activity (SFig. 3e). Due to the variability in timing (SFig. 3f), On events were not evident in traces averaged across the population of ROIs for each cluster (Fig. 2c). Notably, such delayed On events were also observed in some Off-type RGCs (Baden et al., 2016). In contrast, On BCs only rarely exhibited analogous Off responses (SFig. 3e).

A second fundamental principle of inner retinal organisation is the segregation into temporal “transient” and “sustained” channels which map onto the IPL centre and borders, respectively (Awatramani and Slaughter, 2000; Roska and Werblin, 2001; Baden et al., 2012; Borghuis et al., 2013). While our results are broadly in line with this notion, the full picture is more complex: For example, although spike-like events were observed most frequently towards the IPL centre (Fig. 2e,g_3_, SFig. 3g) (Dreosti et al., 2011; Baden et al., 2012; Saszik and DeVries, 2012; Puthussery et al., 2013), they could be found at all IPL depths. In addition, all On but none of the Off cells showed a sustained plateau following an initial fast peak (Fig. 2g_4_), making the most sustained Off cell (C_1_) nearly as transient as the most transient On cell (C_X_, Fig. 2g_5_). Interestingly, a transient or spiking response was not correlated with a short response delay (SFig. 3h,i): For example, C_7_ responses were sustained but had a short delay, whereas C_3a_ responses were transient with a moderate delay. This suggests that temporal properties of BCs like response delay (“fast” vs. “slow” onset) and transience (speed of response decay) can independently vary between BC types - a key ingredient towards the computation of motion (Kim et al., 2014; Serbe et al., 2016).

Finally, we also found a spatial map across the IPL: RF size varied systematically among BC clusters and significantly decreased with increasing stratification depth (Fig. 2f,g_7_; ρ=0.89, p<0.001, n=14 clusters, linear correlation), such that RF diameters of Off BCs (66.3±3.5 μm, mean±s.d.) were on average 10.4 μm larger than those of On BCs (55.9±5.3 μm). Despite this overall trend, RF sizes within one IPL depth differed substantially: For example, C_3b_ or C_5a_ had RF diameters nearly 10 μm larger than C_4_ or C_7_, respectively.

In summary, our results highlight important exceptions from fundamental principles of inner retinal organisation and identify a new spatial organising principle. They indicate that functionally opposite signals such as short and long delays or even On versus Off response polarities co-exist at a single depth.

### BC surround activation increases functional diversity

The organising principles discussed above were extracted from the responses to the local chirp alone, yet responses to the full-field chirp were substantially more heterogeneous (Fig. 2c, Fig. 3). For both On and Off BC clusters, additional surround stimulation significantly decorrelated chirp responses across clusters of the same polarity, and further anticorrelated responses of opposite polarity (Fig. 3a; correlation ρl_oca_l=0.9 vs. ρ_full_-field=0.7 and ρ _loca_l=−0.3 vs. ρfull-field=−0.5, p<0.001, n=14, Wilcoxon signed-rank test). This effect could be quite dramatic (Fig. 3b): For example, Off clusters C_2_ and C_3b_ responded nearly identical to the local chirp, whereas their responses to the full-field chirp were effectively uncorrelated (ρ_loca_l=0.7 vs. ρ_full-fie_ld=0.1). We saw similar differences between On cluster C_6_ and C_9_ responses (ρ _lo_cal=0.9 vs. ρfull-field=0.4). This markedly broadened the response space covered by BC types (Fig. 3c,d,cf. Fig. 2g). For example, during full-field stimulation some On BCs (e.g. C_5a_ or C_9_) lost their sustained plateau phase and became much more transient (Fig. 2c, Fig. 3d_4,5_). The increased diversity is also evident in a two-dimensional representation of the response means for centre-surround relative to centre-only activation (Fig. 3c, Methods). Accordingly, the major type-specific differences in the final output of BCs appear to be determined by concomitant centre and surround activation, rather than centre activation alone.

**Figure 3.**
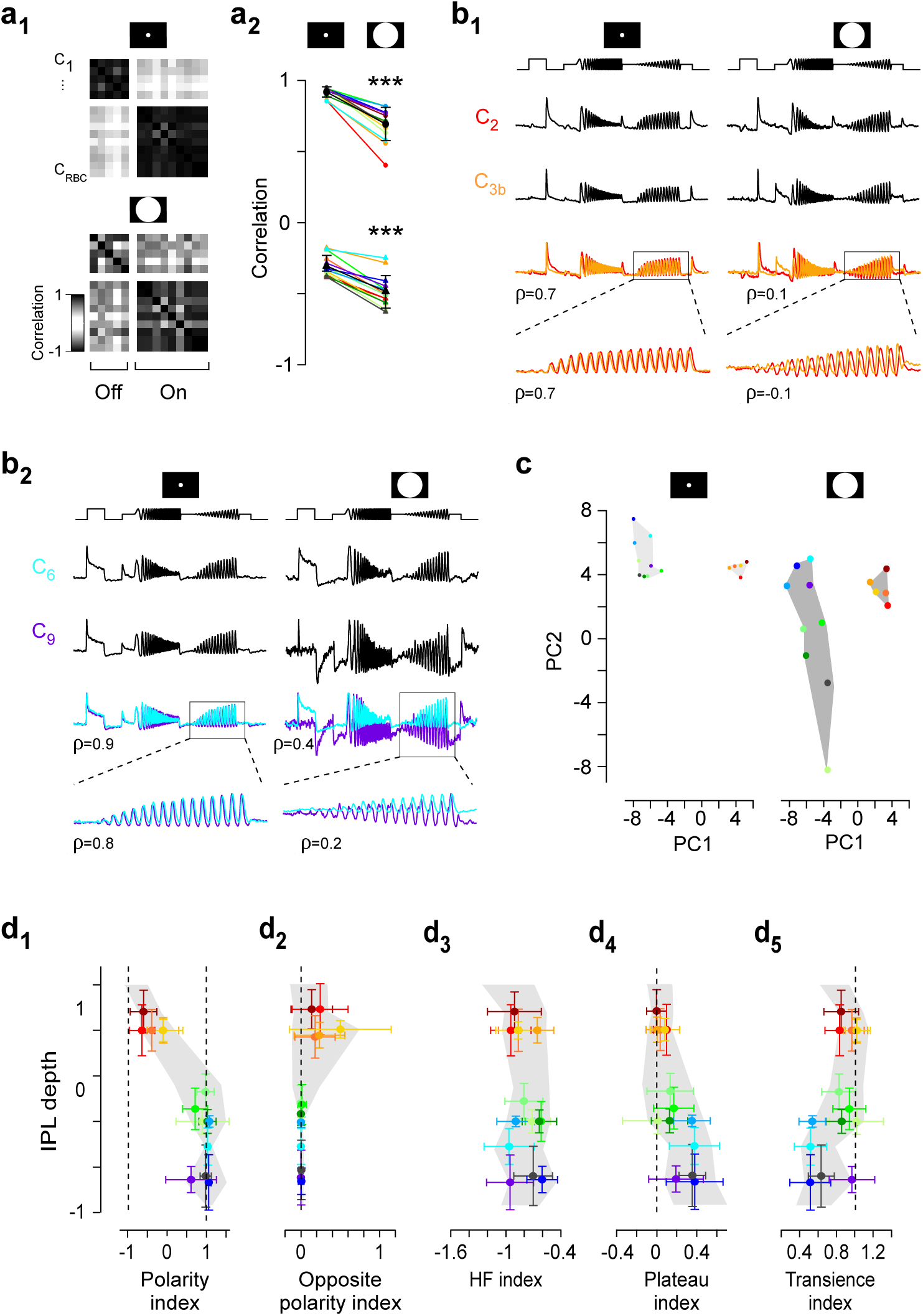
Surround activation increases functional diversity across BCs. **a**, Correlation matrix between cluster means of local (*a_1_, top*) and full-field (*a_1_, bottom*) chirp responses, with darker colours indicating higher (anti)correlation. a_2_, Mean correlation between local and full-field chirp responses for each cluster with all other clusters of the same (*top*) and opposite (*bottom*) polarity (ρl_oca_l =0.9 vs. ρ_fu_l_l-fie_ld=0.7 and ρl_oca_l =−0.3 vs. ρ_fu_ll-field=−0.5, p<0.001, n=14, non-parametric paired Wilcoxon signed-rank test). Mean ± s.d. in black. **b**, Mean chirp responses of two Off (C_2_ and C_3b_, b1) and two On (C_6_ and C_9_, b_2_) clusters, with linear correlation coefficient (ρ) of whole trace or contrast ramp indicated. **c**, Cluster means of local (*left*) and full-field (*right*) chirp responses embedded in two-dimensional feature space based on first and second principal components (PC). **d**, Response measures estimated from full-field chirp responses for all ROIs plotted against IPL depth (cf. Fig. 2g).

### Different ACs mediate and gate BC surround

We next dissected the cellular components underlying the observed surround effects pharmacologically. The two major groups of AC in the mammalian retina, small-and wide-field ACs, use glycine and GABA as their primary neurotransmitter, respectively (Fig. 4a) (Pourcho and Goebel, 1983; Menger et al., 1998). Both groups contact BC axon terminals which in turn express GABA receptors and, in the case of Off BCs, also glycine receptors (Lukasiewicz and Werblin, 1994; Pan and Lipton, 1995; Euler and Wässle, 1998; Euler and Masland, 2000). In addition, there is extensive crosstalk among ACs (Roska et al., 1998; Eggers and Lukasiewicz, 2006; Eggers et al., 2010). To test which of these interactions modulate the BC surround, we pharmacologically blocked either GABA or glycine receptors while monitoring light-evoked glutamate release. Representative for On and Off BCs, we focused these measurements on the BC types overlapping with the On and Off ChAT band, respectively.

**Figure 4.**
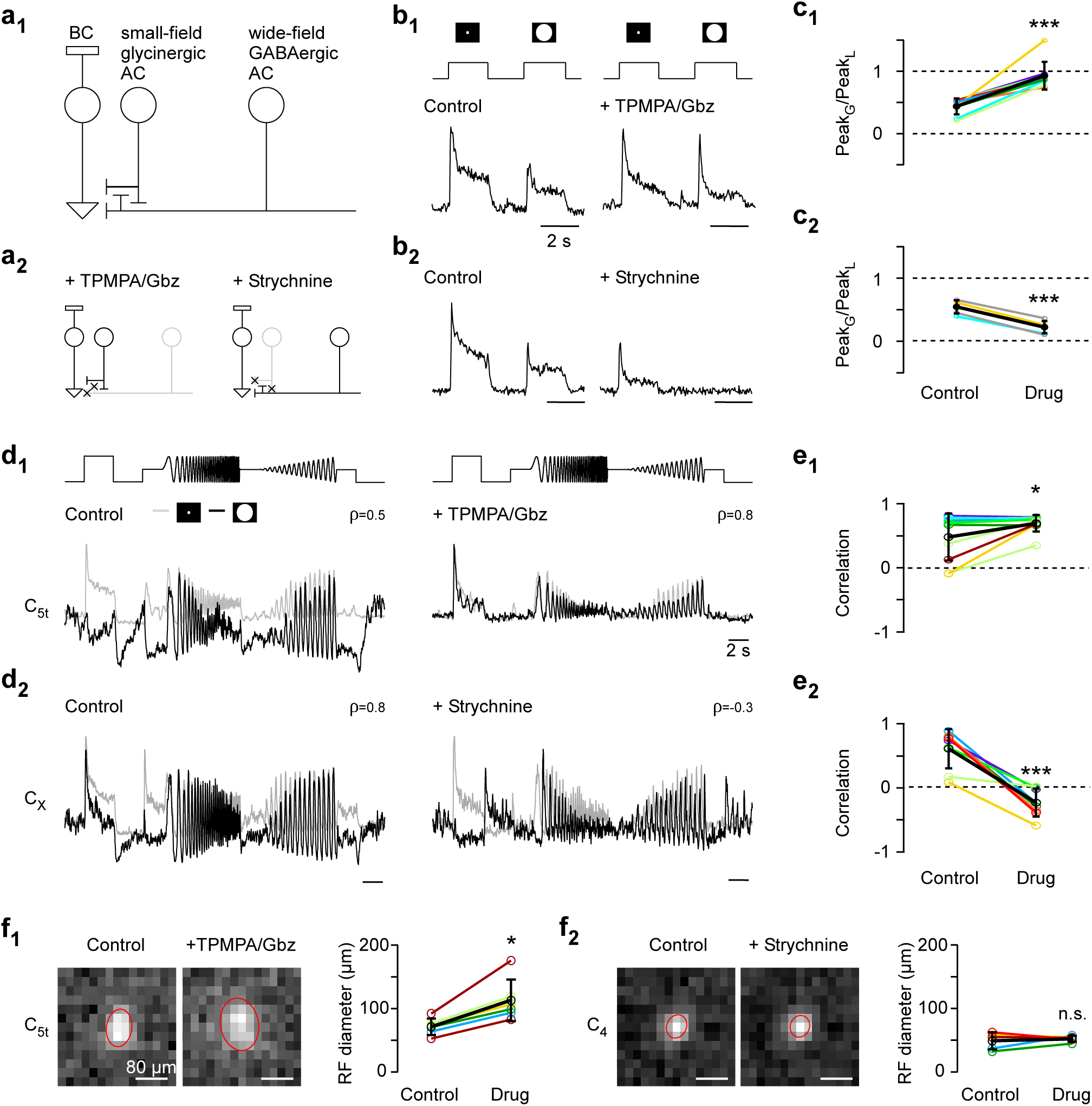
Opposite effects of GABA-and glycinergic ACs on BC output. **a**, Schematic wiring of GABA-and glycinergic ACs with BC terminals (**a**_1_), illustrating the effects of pharmacological GABA (TPMPA/Gbz; 75/10 μM) and glycine (Strychnine; 0.5 μM) receptor block (**a**_2_). **b**, Mean responses (n=5 trials) of individual exemplary ROIs to alternating local and full-field flashes under control conditions and with GABA (b1) or glycine (**b**_2_) receptor block. **c**, Quantification of drug-induced changes in peak response amplitude across different BC clusters upon blocking GABA (**c**_1_; p<0.001, n=9 clusters from 5 scan fields and 4 mice, non-parametric paired Wilcoxon signed-rank test) and glycine receptors (**c**_2_; p<0.001, n=6 clusters from 5 scan fields and 4 mice). Mean ± s.d. in black. For (c-f), traces/RFs were averaged across all ROIs from one scan field assigned to the same cluster (Methods). **d**, Local (*grey*) and full-field (*black*) chirp responses for control and drug conditions (d1: GABA receptor block; d_2_: glycine receptor block), with linear correlation coefficient (ρ) between each pair indicated. **e**, Linear correlation coefficients of local and full-field chirp responses across different clusters for GABA (**e**_1_; p<0.05, n=10 from 5 scan fields and 4 mice, non-parametric paired Wilcoxon signed-rank test) and glycine (**e**_2_; p<0.001, n=8 from 4 scan fields and 3 mice) receptor block. **f**, Spatial RFs with 2 s.d. outline of Gaussian fit shown in red (*left*) and quantification of changes in RF diameter across different BC clusters (*right*) upon blocking GABA (**f**_1_; p<0.05, n=6 cluster from 3 scan fields and 2 mice, non-parametric paired Wilcoxon signed-rank test) and glycine receptors (**f**_2_; p>0.05, n=5 cluster from 3 scan fields and 2 mice).

Pharmacological manipulation had little effect on overall response shape for local stimuli, but caused strong effects for full-field stimulation (Fig. 4). Blocking GABA_A_ and GABA_C_ receptors (with 10 μM Gabazine (Gbz) and 75 μM TPMPA, respectively) lead to an increase in response amplitude in both On and Off BCs (Fig. 4b_1_,c_1_, SFig. 4a; ρ_Control_=0.44 vs. ρ_Drug_=0.93, p<0.001, n=9 clusters), consistent with attenuated surround inhibition (Roska et al., 2000; Ichinose and Lukasiewicz, 2005; Buldyrev and Taylor, 2013). In addition, blocking GABA receptors nearly eliminated the difference between local and full-field chirp responses (Fig. 4d_1_,e_1_, SFig. 4b; ρ_Control_=0.58 vs. ρ_D_ru_g_=0.8, p<0.05, n=10 clusters). This suggests that the BC surround is largely generated by presynaptic inhibition from GABAergic ACs (Discussion).

In contrast, blocking glycine receptors (with 0.5 μM Strychnine) reduced the response to full-field flashes (Fig. 4b_2_,c_2_, SFig. 4a; ρ_Control_=0.55 vs ρ_Drug_=0.22, p<0.001, n=6 clusters), consistent with an increase in surround strength. Notably, this effect reliably induced a polarity switch in BC responses to full-field stimulation, thus anti-correlating local and full-field chirp responses (Fig. 4d,e_2_, SFig. 4c; ρ_Control_=0.6 vs. ρ_Drug_=−0.24, p<0.001, n=8 clusters). This suggests that glycinergic ACs primarily modulate BC output in an indirect way by inhibiting GABAergic ACs, leading to decreased inhibition. In the absence of drugs, a polarity switch could also be consistently induced by presenting an annulus chirp that excluded the central 100 μm-spot (SFig. 4d) (Dacey et al., 2000). Accordingly, the BC surround not only modulates an existing centre response, but can also act as an independent “input system” that by itself is capable of driving glutamate release from BCs. One explanation is that inhibition modulates a light-independent tonic release of glutamate, which appears to be a common feature of BCs (Venkataramani and Taylor, 2016).

Additionally, we found that blocking GABA receptors increased the size of the BC RF centre, whereas glycine receptor block had no detectable effect on RF size (Fig. 4f). This implies that not only temporal, but also spatial properties of the BC centre depend on the state of the GABAergic inhibitory network in the inner retina.

In summary, the two major groups of ACs of the mouse retina appear to act in tandem to set the balance of excitation and inhibition, thereby increasing functional diversity among BC types.

### Differential centre-surround interactions underlie BC diversity

An increase in functional diversity upon surround stimulation is only possible if surround networks of different BC types differentially process visual stimuli. Accordingly, it is important to obtain precise estimates not only of centre but also of surround spatio-temporal RFs. To this end, we estimated a series of linear time kernels at different distances from the BC RF centre by presenting independently flickering narrow concentric rings (“ring noise”, Fig. 5a) and isolated RF centre and surround space and time kernels for every BC cluster (Fig. 5b,c, SFig. 5a-c, Methods).

**Figure 5.**
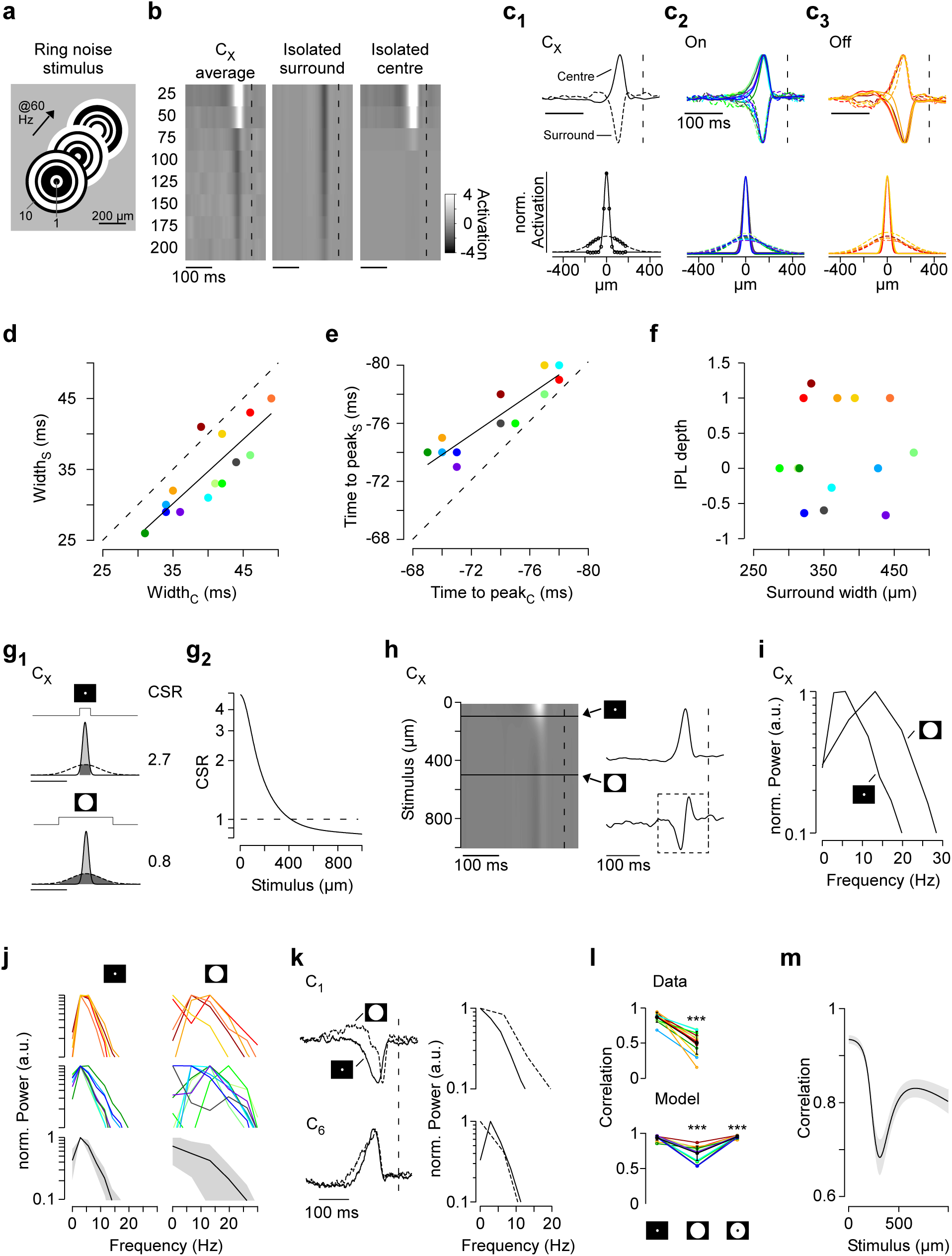
Differential centre-surround organisation underlies BC functional diversity. **a**, Schematic of ring noise stimulus, showing intensity distribution across rings (width: 25 μm) for the first three stimulus frames. **b**, Centre-surround maps of an example cluster (**C**_X_), depicting temporal kernels of n=8 rings with increasing distance from the scan field centre. From the average map of each cluster (*left*), surround (*middle*) and centre (*right*) components were isolated using Singular value decomposition (SVD; Methods, cf. SFig. 5a). Dashed lines at t = 0. **c**, Normalised time (*top*) and space (*bottom*) kernels of C_X_ (**c**_1_) and all On (**c**_2_) and Off (**c**_3_) BC clusters. Space kernels represent extrapolated Gauss fits of centre and surround activation across rings shown in (b), with circles corresponding to the original data points (Methods). **d**, Half-maximal width was consistently narrower for centre compared to surround time-kernels (r=0.83, p<0.001, n=14, linear correlation; slope=0.93, p>0.05, linear regression for slope=1). Black line corresponds to linear fit and for the dashed line the slope=1. **e**, Time to peak of centre time kernels preceded peak of surround time kernels (r=0.91, p<0.001, n=14, linear correlation). This effect was stronger for cells with shorter time to peak (slope=0.65, p<0.001, linear regression for slope=1). **f**, Half-maximal width of surround space kernels did not correlate with mean cluster IPL depth (r=0.06, p>0.05, n=14, linear correlation). **g**, Space kernels of C_X_ (g1), with predicted centre-surround activation ratios (CSR) of a local (*top*) and full-field (*bottom*) spot stimulus obtained from the relative activation of centre (*light grey*) and surround (*dark grey*). g_2_, Predicted CSRs of C_X_ for different spot diameters. **h**, Normalised effective time kernels of C_X_ for different stimulus diameters (*left*), obtained from a weighted addition of centre and surround time kernels shown in (c_1_). *Right*, Normalised effective time kernels estimated for a stimulus diameter of 100 and 500 μm, respectively. Spectra in (i) and (j) were estimated from kernel fraction as indicated by dashed rectangle. **i**, Predicted normalised spectra estimated from effective time kernels shown in (h, *right*). **j**, Predicted normalised spectra of Off (*top*) and On (*middle*) BC clusters during local (*left*) and full-field (*right*) stimulation. Average spectra across BC cluster and s.d. shown in black and grey, respectively (*bottom*). **k**, Measured, normalised time kernels (*left*) and normalised frequency spectra (*right*) estimated for a local and full-field spot noise stimulus, exemplary shown for C_1_ and C_6_. **l**, Correlation of time kernels estimated from the spot noise (p<0.001, n=13, non-parametric paired Wilcoxon signed-rank test) and predicted by the model (for both p<0.001, n=14) for local, full-field and surround-only stimulation. Mean ± s.d. in black. **m**, Average correlation across predicted cluster time kernels for different stimulus diameters. Errors in s.e.m.

Centre and surround time kernels of the same clusters were correlated with respect to width and time to peak (Fig. 5d,e). Centre kernels were consistently broader than surround kernels (Fig. 5d; 5.3±3.2 ms, mean±s.d.), while the time lag between centre and surround peaks was more pronounced for cells with shorter time to peak for both On and Off clusters (Fig. 5e) with the delay ranging from 1 to 5 ms. This indicates that BC types may not contribute strongly to generating their own surround; if this had been the case, one should have observed a slope of 1 (Fig. 5e). One possible mechanism is that ACs providing surround inhibition to BCs with short delays are driven by BCs with longer delays. Moreover, the spatial extent of surround kernels varied between types (Fig. 5f, range: 287-478 μm), indicating that also different AC networks are involved in different BC circuits.

Next, we investigated how functional diversity may emerge from the observed differences in centre-surround organisation. Using the spatial RFs, we first estimated the centre-surround ratio (CSR) for circular stimuli of different sizes for each cluster (Fig. 5g, SFig. 5d, Methods). For an example cluster (C_X_), a small stimulus (100 μm in diameter) resulted in a two-fold stronger centre activation compared to the surround (CSR=2.3, “centre-dominant”). In contrast, the surround was stronger than the centre for a full-field spot (500 μm in diameter, CSR=0.8, “surround-dominant”). Although the exact CSRs slightly differed between BC clusters for a given stimulus size, all clusters gradually switched from a centre-dominant to a surround-dominant mode of operation between 200 and 600 μm-diameter stimuli (SFig. 5d).

We next used a simple model to predict how the temporal properties of BCs change with the spatial extent of the stimulus (Fig. 5h). For example, the predicted kernel of C_X_ “accelerated” with stimulus size, approximately doubling its centre frequency and temporal bandwidth (Fig. 5i). The model predicted cluster-specific changes in the temporal coding properties with increasing stimulus size, leading to an increase in the overall diversity across clusters (Fig. 5j). As a result, combined kernels for larger stimuli encompassed a much broader range of temporal frequencies (Fig. 5i). To directly test this prediction experimentally, we recorded BC responses to flickering spots at two different stimulus sizes (100 and 500 μm diameter, respectively). In agreement with the model, time kernels consistently differed between stimulus sizes (Fig. 5k), leading to lower time kernel correlations across clusters for centre-surround compared to centre-or surround-only stimulation (Fig. 5l, cf. Fig. 3a, SFig. 5e). In addition, the model predicts that this effect was strongest between ~200 and 500 μm (Fig. 5m), matching the distribution of RF centre sizes of RGCs in the mouse retina (Baden et al. 2016).

## DISCUSSION

We systematically surveyed the visual response properties of mouse BCs by imaging their glutamatergic output and arrive at a census of the excitatory signals that drive inner retinal circuits. Beyond the expected mapping of “classical” functional BC properties like On and Off or different response kinetics onto specific strata of the IPL (Awatramani and Slaughter, 2000; Roska and Werblin, 2001; Baden et al., 2012), we found that the overall functional heterogeneity within individual IPL strata was larger than previously thought. We showed how this temporal diversity is created by the interplay of the excitatory drive forwarded from the dendrites and local axonal inputs from ACs, with the two input streams acting at different spatial scales. As a result, the spatial structure of the visual input fundamentally sets the balance between excitation and inhibition and thus the temporal encoding in BCs – and consequently the visual system.

### Sources of glutamate release in the IPL

One central assumption was that BC axon terminals are the source of glutamate release in the IPL. However, some On BCs feature ectopic synapses along their axon (Dumitrescu et al., 2009; Kim et al., 2012) which might contribute to observed On responses in the Off sub-lamina (Fig. 2d,g_2_). Two findings strongly argue against this possibility: First, On responses in the Off sub-lamina were always accompanied by a clear and dominant Off component (Fig. 2d,g_1_). Second, they were delayed relative to On responses in the On layer (SFig. 3f), suggesting a different origin. Instead, they likely either resulted from surround inhibition or from intrinsic properties of some Off BC types, such as a depolarising rebound following transient inhibition (Mitra and Miller, 2007a, 2007b).

In addition, the mouse retina harbours at least one AC type that uses glutamate as neurotransmitter at conventional non-ribbon synapses (Lee et al., 2014). This glutamatergic AC (GAC) stratifies between On and Off ChAT bands and receives inputs from both On and Off BCs (Lee et al., 2014). Dendritic calcium imaging suggests that GACs process BC input locally, resulting in On and Off responses in the On and Off sub-lamina, respectively (Lee et al., 2015). These signals are expected to be very similar to the respective BC inputs and can likely not be separated using our clustering method. Therefore, we took the conservative approach and did not include the GAC in our analysis.

In contrast, we included RBCs as they displayed robust light-evoked calcium responses at our stimulus intensities in the low-photopic regime (Fig. S2e,f). Under these conditions, rod photoreceptors, which provide the excitatory synaptic input to RBCs (Dacheux and Raviola, 1986; Bloomfield and Dacheux, 2001; Keeley and Reese, 2010), are thought to be saturated. However, recent evidence suggests that both rods (Tikidji-Hamburyan et al.,2014) and RBCs (Chen et al., 2014b) can be active under photopic conditions. Perhaps direct contacts between RBCs and cones identified at both the ultrastructural (Behrens, Schubert, Haverkamp, Euler and Berens, personal communication) and functional level (Pang et al., 2010) contribute to the observed responses, further challenging the view that RBCs solely mediate vision in dim light.

### The origin of BC functional diversity

We found that functional diversity amongst BCs is primarily driven by a change in the balance of excitation and GABAergic inhibition, which in turn is set by glycinergic inhibition. Specifically, GABAergic inhibition seems to be distinct for different BC types and can thus act to decorrelate BC channels (Figs. 3-5).

The decorrelating surround effects are likely due to AC mediated inhibition in the inner retina, rather than horizontal cell (HC) mediated inhibition in the outer retina: First, mouse HCs (Peichl and González-Soriano, 1994) have little effect on BC (Purgert and Lukasiewicz,2015) and RGC function (Cook and McReynolds, 1998; Taylor, 1999; Ichinose and Lukasiewicz, 2005). Second, we found that the GABAergic BC surround was gated by glycinergic signals (Fig. 4) which appear to be exclusive to the inner retina (Wässle et al., 2009a).

Our results suggest that in the intact (whole-mounted) retina, glycinergic effects via serial pathways – such as the gating of GABAergic inhibition to BCs – predominate over direct inputs to BCs. While such serial inputs exist in salamander (Zhang et al., 1997; Roska et al., 2000), available data for the mouse is less conclusive (Eggers and Lukasiewicz, 2006; Majumdar et al., 2009). In mouse, glycinergic ACs were found to mainly mediate vertical signalling (crossover inhibition) via direct inputs to Off BCs (Ivanova et al., 2006; Roska et al., 2006; Chavez and Diamond, 2008; Eggers and Lukasiewicz, 2011). However, these studies were performed in vertical slices where lateral connections are likely compromised.

### The elementary computational unit of the IPL

The powerful effect of inner retinal inhibition on visual encoding in BCs bolsters the view that the elementary computational unit of the IPL must be the individual BC synaptic terminal (Eggers and Lukasiewicz, 2011; Oesch and Diamond, 2011; Odermatt et al., 2012; Baden and Euler, 2013; Nikolaev et al., 2013). Clearly, BCs are not electrotonically compact units, where the computational output can be assessed equally in each compartment (Oltedal et al., 2006). Instead it is at the terminals where two input streams of comparable strength combine: one feedforward pathway reflecting the integrated dendritic drive from the photoreceptors and one indirect pathway reflecting the aggregate inhibitory AC network activity. Critically, each input stream by itself can modulate or even drive BC synaptic release.

This further raises the possibility that individual terminals of a single BC may signal independently, if they receive differential inputs from ACs. To what extent such heterogeneity at the level of the BC axon terminal (“presynaptic multiplexing”) matters is actively debated (Euler et al., 2014). While direct evidence for heterogeneity at the level of mouse BC terminal calcium is lacking (Fig. S2b) (Yonehara et al., 2013; Chen et al., 2014b), electrical recordings in salamander retina showed that individual BCs can elicit different responses in different postsynaptic RGCs (Asari and Meister, 2012, 2014). Although this effect could be explained by presynaptic heterogeneity generated by differential AC inputs or by differences in terminal size (Baden et al., 2014), it could also originate from selective postsynaptic inhibition (Masland, 2012b). Additional diversity could come from type-specific differences in the dendritic properties of postsynaptic neurons (Marc and Jones, 2002; Sun et al., 2003; Oesch et al., 2005). For our recording conditions, type-specific functional differences across mouse BCs appear to be more pronounced than any putative differences across terminals of a single cell.

### Decomposition of the visual signal in the inner retina

Why does the mouse retina split the visual signal into 14 parallel channels at the level of BCs? The finding that small stimuli in the range of a BCs dendritic field evoke highly correlated responses among types of the same response polarity implies that the set of BC types is not optimised to decompose the visual signal at this scale. Instead, it is only upon spatially extended stimulation that the rich functional diversity among BC types is revealed.

Our data suggests that this diversity is generated by the interaction of correlated yet not-identical pairs of temporal BC centre and surround kernels. Differences between temporal RFs of centre and surround are emphasised as increasing stimulus size shifts the balance between centre-and surround-activation towards a stronger surround contribution. The outcome is a new “mixed” temporal RF that is – up to a certain stimulus diameter – increasingly distinct from similarly “mixed” temporal RFs of other centre-surround pairs (Fig. 5). Interestingly, the stimulus size leading to maximal diversity matches the average centre size of RGC RFs (Baden et al., 2016). This also implies that a hypothetical RGC type driven by only one BC type should receive temporally distinct inputs from its presynaptic partners depending on the spatial aspects of the stimulus (cf. (Schwartz et al., 2012)).

The notion that an antagonistic centre-surround RF organisation can decorrelate neural response properties is a fundamental principle in neuroscience (e.g. (Barlow, 1961; Atick and Redlich, 1990, 1993)). For example, surround-mediated decorrelation of neurons has been demonstrated for RGCs (Pitkow and Meister, 2012), neurons in primary visual cortex (Vinje et al., 2007) as well as for neurons in other sensory systems (Wiechert et al., 2010). However, these studies focused on decorrelation between neurons independent of cell type. In contrast, less is known about the effect of surround inhibition in decreasing the redundancy of the encoding in different neuronal cell types of the same class. Here, we show that the balance between feedforward excitation and lateral inhibition in the IPL decorrelates parallel signal channels formed by BC types. Such decorrelation is typically linked to efficient coding (Denève and Machens, 2016) of visual stimuli and our data place this computation already at the second synapse of the mouse visual system.

## METHODS

### Animals and tissue preparation

All animal procedures adhered to the laws governing animal experimentation issued by the German Government. For all experiments, we used 3- to 12-week-old Chat^tm2(cre)Lowl^ (n=29; “ChAT:Cre”, JAX 006410, The Jackson Laboratory, Bar Harbor, US) and Tg(Pcp2-cre)1Amc (n=5; “Pcp2”, JAX 006207) mice of either sex. These transgenic lines were cross-bred with the Cre-dependent red fluorescence reporter line Gt(ROSA)26Sor^tm9(CAG-tdTomato)Hze^ (“Ai9^tdTomato^”, JAX 007905) for a subset of experiments.

Animals were housed under a standard 12 hr day/night rhythm. For recordings, animals were dark-adapted for ≥1 h, then anesthetised with isoflurane (Baxter, Unterschleißheim, Germany) and killed by cervical dislocation. The eyes were enucleated and hemisected in carboxygenated (95% O_2_, 5% CO_2_) artificial cerebral spinal fluid (ACSF) solution containing (in mM): 125 NaCl, 2.5 KCl, 2 CaCl_2_, 1 MgCl_2_, 1.25 NaH_2_PO_4_, 26 NaHCO_3_, 20 glucose, and 0.5 L-glutamine (pH 7.4). Then, the tissue was moved to the recording chamber of the microscope, where it was continuously perfused with carboxygenated ACSF at ~37°C. The ACSF contained ~0.1 μM Sulforhodamine-101 (SR101, Invitrogen, Darmstadt, Germany) to reveal blood vessels and any damaged cells in the red fluorescence channel. All procedures were carried out under very dim red (>650 nm) light.

### Virus injection

A volume of 1 μl of the viral construct (AAV9.hSyn.iGluSnFr.WPRE.SV40 or AAV9.CAG.Flex.iGluSnFr.WPRE.SV40 (referred to as “AAV9.iGluSnFr”) or AAV9.Syn.Flex.GCaMP6f.WPRE.SV40, Penn Vector Core, Philadelphia, USA) was injected into the vitreous humour of 3- to 8-week-old mice anesthetised with 10% ketamine (bela-pharm GmbH & Co. KG, Vechta, Germany) and 2% xylazine (Rompun, Bayer Vital GmbH, Leverkusen, Germany) in 0.9% NaCl (Fresenius, Bad Homburg, Germany). For the injections, we used a micromanipulator (World Precision Instruments, Sarasota, USA) and a Hamilton injection system (syringe: 7634-01, needles: 207434, point style 3, length 51 mm, Hamilton Messtechnik GmbH, Hoechst, Germany). Due to the fixed angle of the injection needle (15°), the virus was applied to the ventro-nasal retina. Imaging experiments were performed 3 to 4 weeks after injection.

### Pharmacology

All drugs were bath applied for at least ten minutes before recordings. The following drug concentrations were used (in μM): 10 Gabazine (Tocris Bioscience, Bristol, UK) (Kemmler et al., 2014), 75 TPMPA (1,2,5,6-Tetrahydropyridin-4-yl)methylphosphinic acid, Tocris Bioscience) (Kemmler et al., 2014), and 0.5 strychnine (Sigma-Aldrich, Steinheim am Albuch, Germany) (Schubert et al., 2008). Drug solutions were carboxygenated and warmed to ~37°C before application. Pharmacological experiments were exclusively performed in the On and Off ChAT-immunoreactive bands, which are labelled in red fluorescence in ChAT:Cre x AI9^tdTomato^ crossbred animals.

### Two-photon imaging and light stimulation

We used a MOM-type two-photon microscope (designed by W. Denk, MPI, Heidelberg; purchased from Sutter Instruments/Science Products, Hofheim, Germany). Design and procedures were described previously (Euler et al., 2009). In brief, the system was equipped with a mode-locked Ti:Sapphire laser (MaiTai-HP DeepSee, Newport Spectra-Physics, Darmstadt, Germany), two fluorescence detection channels for iGluSnFR or GCaMP6f (HQ 510/84, AHF/Chroma Tübingen, Germany) and SR101/tdTomato (HQ 630/60, AHF), and a water immersion objective (W Plan-Apochromat 20x/1,0 DIC M27, Zeiss, Oberkochen, Germany). The laser was tuned to 927 nm for imaging iGluSnFr, GCaMP6f or SR101, and to 1,000 nm for imaging tdTomato. For image acquisition, we used custom-made software (ScanM, by M. Müller, MPI, Martinsried, and T.E.) running under IGOR Pro 6.3 for Windows (Wavemetrics, Lake Oswego, OR, USA), taking time-lapsed 64 × 16 pixel image scans (at 31.25 Hz) for glutamate and 32 × 32 pixel image scans (at 15.625 Hz) for calcium imaging. For visualising morphology, 512 × 512 pixel images were acquired.

For light stimulation, we focused a DLP projector (K11, Acer) through the objective, fitted with band-pass-filtered light-emitting diodes (LEDs) (“green”, 578 BP 10; and “blue”, HC 405 BP 10, AHF/Croma) to match the spectral sensitivity of mouse M-and S-opsins. LEDs were synchronised with the microscope’s scan retrace. Stimulator intensity (as photoisomerisation rate, 10^3^ P^*^/s/cone) was calibrated as described previously (Euler et al., 2009) to range from 0.6 and 0.7 (black image) to 18.8 and 20.3 for M-and S-opsins, respectively. Due to technical limitations, intensity modulations were weakly rectified below 20% brightness. An additional, steady illumination component of ~10^4^ P^*^/s/cone was present during the recordings because of two-photon excitation of photopigments (for detailed discussion, see (Euler et al., 2009) and (Baden et al., 2013)). The light stimulus was centred prior to every experiment, such that its centre corresponded to the centre of the recording field. For all experiments, the tissue was kept at a constant mean stimulator intensity level for at least 15 s after the laser scanning started and before light stimuli were presented. Four types of light stimuli were used (Fig. 1): (*i*) Full-field (800x600 μm) and (*ii*) local (100 μm in diameter) “chirp” stimuli consisting of a bright step and two sinusoidal intensity modulations, one with increasing frequency (0.5-8 Hz) and one with increasing contrast, (*iii*) 1 Hz light flashes (500 μm in diameter, 50% duty cycle), and (*iv*) binary dense noise (20×15 matrix of 20×20 μm pixels; each pixel displayed an independent, balanced random sequence at 5 Hz for 5 minutes) for space-time receptive field (RF) mapping. In a subset of experiments, we used two additional stimuli: (*v*) A “ring noise” stimulus (10 annuli with increasing diameter, each annulus 25 μm wide), with each ring’s intensity determined independently by a balanced 68 s random sequence at 60 Hz repeated four times, and (*vi*) a surround chirp stimulus (annulus; Full-field chirp sparing the central 100 μm corresponding to the local chirp). For all drug experiments, we showed in addition (*vii*) a stimulus consisting of alternating 2 s full-field and local light flashes (500 and 100 μm in diameter, respectively). All stimuli were achromatic, with matched photo-isomerisation rates for mouse M-and S-opsins.

### Estimating recording depth in the IPL

For each scan field, we used the relative positions of the inner (ganglion cell layer) and outer (inner nuclear layer) blood vessel plexus to estimate IPL depth. To relate these blood vessel plexi to the ChAT bands, we performed separate experiments in ChAT:Cre x AI9^tdTomato^ mice: High resolution stacks throughout the inner retina were recorded in the ventro-nasal retina. The stacks were then first corrected for warping of the I PL using custom-written scripts in IGOR Pro. In brief, a raster of markers (7 × 7) was projected in the x-y plane of the stack and for each marker the z positions of the On ChAT band was manually determined. The point raster was used to calculate a smoothed surface, which provided a z offset correction for each pixel beam in the stack. For each corrected stack, the z profiles of tdTomato and SR101 labelling were extracted by manually drawing ROIs in regions where only blood vessel plexi or the ChAT bands were visible. The two profiles were then matched such that 0 corresponded to the inner vessel peak and 1 corresponded to the outer vessel peak. We averaged the profiles of n=9 stacks from 3 mice and determined the I PL depth of On and Off ChAT band to be 0.48 ± 0.011 and 0.77 ± 0.014 (mean ± s.d.), respectively. The s.d. corresponds to an error of 0.45 and 0.63 μm for On and Off ChAT band, respectively. In the following, recording depths relative to blood vessel plexi were transformed into I PL depths relative to ChAT bands for all scan fields (Fig. 1c), with 0 corresponding to the On ChAT band and 1 corresponding to the Off ChAT band.

### Data analysis

Data analysis was performed using Matlab 2015a (The Mathworks Inc., Ismaning, Germany), and IGOR Pro. Data were organised in a custom written schema using the *DataJoint for Matlab* framework (github.com/datajoint/datajoint-matlab) (Yatsenko et al., 2015). All data as well as Matlab code including basic visualisation routines are available at www.retinal-functomics.org.

#### Pre-processing

Regions-of-interest (ROIs) were defined automatically by a custom correlation-based algorithm in IGOR Pro. For this, the activity stack in response to the dense noise stimulus (64 × 16 × 10,000 pixels) was first de-trended by high-pass filtering the trace of each individual pixel above ~0.1 Hz. For the 100 best responding pixels in each recording field (highest s.d. over time), the trace of each pixel was correlated with the trace of every other pixel in the field. Then, the correlation coefficient (ρ) was plotted against the distance of the two pixels and the average across ROIs was computed (SFig. 1a1). A scan field-specific correlation threshold (*ρ*_*Threshold*_) was determined by fitting an exponential between the smallest distance and 5 μm (SFig. 1a1). *ρ*_*Threshold*_ was defined as the correlation coefficient at x = λ, where λ is the exponential decay constant (space constant; SFig. 1a_2_). Next, we grouped neighbouring pixels with **ρ** > *ρ*_*Threshold*_ into one ROI (SFig. 1b). To match ROI sizes with the size of BC axon terminals, we restricted ROI diameters (estimated as effective diameter of area-equivalent circle) to range between 0.75 and 4 μm (SFig. 1a_2_,d). For validation, the number of ROIs covering single axon terminals was quantified manually for n=31 terminals from n=5 GCaMP6-expressing BCs (SFig. 1d, 2a).

The glutamate (or calcium) traces for each ROI were extracted (as ΔF/F) using the image analysis toolbox SARFIA for IGOR Pro (Dorostkar et al., 2010). A stimulus time marker embedded in the recorded data served to align the traces relative to the visual stimulus with 2 ms precision. For this, the timing for each ROI was corrected for sub-frame time-offsets related to the scanning. Stimulus-aligned traces for each ROI were imported into Matlab for further analysis.

For the chirp and step stimuli, we then subtracted the baseline (median of first 20-64 samples), computed the median activity *r(t)* across stimulus repetitions (5 repetitions for chirp, 40-50 repetitions for step) and normalised it such that max_t_(∣(t)∣) = 1.

#### Receptive field mapping

We mapped the linear RFs of the neurons by computing the glutamate/calcium transient-triggered average. To this end, we resampled the temporal derivative of the glutamate/calcium response ċ(t) at 10-times the stimulus frequency and used Matlab’s findpeaks function to detect the times t_i_ at which transients occurred. We set the minimum peak height to 1 s.d., where the s.d. was robustly estimated using:

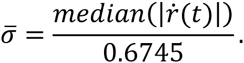

We then computed the glutamate/calcium transient-triggered average stimulus, weighting each sample by the steepness of the transient:

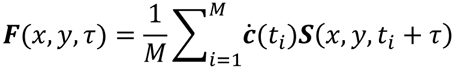

Here, ***S***(*x,y,t*) is the stimulus, *τ* is the time lag (ranging from approx. −297 to 1,267 ms) and *M* is the number of glutamate/calcium events. We smoothed this raw RF estimate using a 3×3 pixel Gaussian window for each time lag separately. We used singular value decomposition (SVD) to extract temporal and spatial RF kernels.

To extract the RF’s position and scale, we fitted it with a 2D Gaussian function using Matlab’s lsqcurvefit. RF quality *(RF_qi_)* was measured as one minus the fraction of residual variance not explained by the Gaussian fit
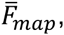

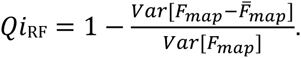

#### Other response measures

<u>Response quality index</u> – To measure how well a cell responded to a stimulus (local and fullfield chirp, flashes), we computed the signal-to-noise ratio

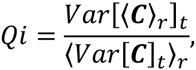

where ***C*** is the *T* by *R* response matrix (time samples by stimulus repetitions) and 〈 〉_x_ and *Var*[ ]_x_ denote the mean and variance across the indicated dimension, respectively (Baden et al., 2016).

For further analysis, we used only cells that responded well to the local chirp stimulus (*Qi*_lchirp_>0.3) and resulted in good RFs (*Qi*_RF_ > 0.2).

<u>Polarity index</u>- To distinguish between On and Off BCs, we calculated the Polarity index *(POi)* from the step response to local and full-field chirp, respectively, as

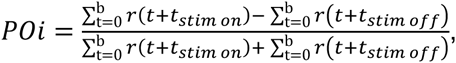

where *b* = 2 s (62 samples). For cells responding solely during the On-phase of a step of light *POi* = 1, while for cells only responding during the step’s Off-phase *POi* = −1.

<u>Opposite polarity index</u> – The number of opposite polarity events (*OPi*) was estimated from individual trials of local and full-field chirp step responses (first 6 seconds) using IGOR Pro’s FindPeak function. Specifically, we counted the number of events that occurred during the first 2 seconds following the step onset and offset for Off and On BCs, respectively. For each trial the total number of events was divided by the number of stimulus trials. If *OPi* = 1, there was on average one opposite polarity event per trial.

<u>High frequency index</u> – The high frequency index (*HFi*) was used to quantify spiking (cf. (Baden et al., 2012)) and was calculated from responses to individual trials of the local and full-field chirp, respectively. For the first 6 seconds of each trial, the frequency spectrum was calculated by fast Fourier transform (FFT) and spectra were averaged across trials for individual ROIs. Then, *HFi* = log(F_1_/F_2_), where F_1_ and F_2_ are the mean power between 0.5-1 Hz and 2-16 Hz, respectively.

<u>Response transience index</u> – The step response (first 6 seconds) of local and full-field chirps was used to calculate the response transience (*RTi*). Traces were up-sampled to 500 Hz and the response transience was calculated as

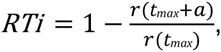

where *α*= 400 ms is the read-out time following the peak response *t*_*max*_. For a transient cell with complete decay back to baseline *RTi* =1, whereas for a sustained cell with no decay *RTi* =0.

<u>Response plateau index</u> – Local and full-field chirp responses were up-sampled to 500 Hz and the plateau index (*RPi*) was determined as:

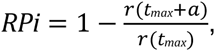

with the read-out time *α*= 2 s. A cell showing a sustained plateau has an *RPi* =1, while for a transient cell *RPi* =0.

<u>Response delay</u> – The response delay (*t*_*delay*_) was defined as the time from stimulus onset/offset until response onset and was calculated from the up-sampled local chirp step response. Response onset (*t*_*onset*_) and delay (*t*_*delay*_) were defined as

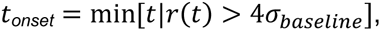

and

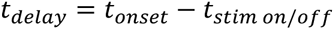

respectively.

#### Feature extraction

We used sparse principal component analysis, as implemented in the SpaSM toolbox by (Sjöstrand and Clemmensen, 2012), to extract sparse response features from the mean responses across trials to the local (12 features) and full-field chirp (6 features), and the step stimulus (6 features) (as described in (Baden et al., 2016), SFig. 3a). Before clustering, we standardised each feature separately across the population of cells.

#### Anatomy-guided clustering

BC-terminal volume profiles were obtained from EM-reconstructions of the inner retina (Greene et al., n.d.; Kim et al., 2014) To isolate synaptic terminals, we removed those parts of the volume profiles which likely correspond to axons. We estimated the median axon density for each type from the upper 0.06 units of I PL and subtracted twice that estimate from the profiles, clipping at zero. Profiles were smoothed with a Gaussian kernel (s.d.=0.14 units I PL depth) to account for jitter in depth measurements of two-photon data.

We used a modified Mixture of Gaussian model (Szczurek et al., 2010) to incorporate the prior knowledge from the anatomical BC profiles. For each ROI *i* with I PL depth d_i_, we define a prior over anatomical types *c* as

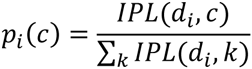

where *IPL*(d,c) is the IPL terminal density profile as a function of depth and anatomical cell type. The parameters of the Mixture of Gaussian model are estimated as usual, with the exception of estimating the posterior over clusters. Here, the mixing coefficients are replaced by the prior over anatomical types, resulting in a modified update formula for the posterior:

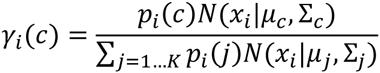

All other updates remain the same as for the standard Mixture of Gaussians algorithm (Bishop, 2006). We constrained the covariance matrix for each component to be diagonal, resulting in 48 parameters per component (24 for the mean, 24 for the variances). We further regularised the covariance matrix by adding a constant (10^−5^) to the diagonal.

For each pair of clusters, we computed the direction in feature space that optimally separated the clusters

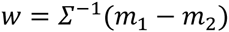

where *m*_i_ are the cluster means in feature space and *Σ* is the pooled covariance matrix. We projected all data on this axis and standardised the projected data according to cluster 1 (i.e. subtract the projected mean of cluster 1 and divide by its s.d.). We compute *d’*as a measure of the separation between the clusters:

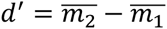

where 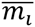

are the means of the two clusters in the projected, normalised space.

#### Further statistical analysis

<u>Field entropy</u> - Field entropy (*S*_*Field*_) was used as a measure of cluster heterogeneity within single recording fields and was defined as

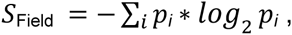

where *i* is the number of clusters in one recording field and *p*_*i*_ corresponds to the number of ROIs assigned to the *i*^th^ cluster. *S*_*Field*_ = 0 if all ROIs of one recording field are assigned to one cluster and *S*_*Field*_ increases if ROIs are equally distributed across multiple clusters. In general, high field-entropy indicates high cluster heterogeneity within a single field.

<u>Analysis of response diversity</u> – To investigate the similarity of local and full-field chirp responses across clusters (Fig. 3), we determined the linear correlation coefficient between any two cluster pairs. The analysis was performed on cluster means. For every cluster, correlation coefficients were averaged across clusters with the same and opposite response polarity, respectively. We used principal component analysis (using Matlab’s pca function) to obtain a 2D embedding of the mean cluster responses. The PCA was computed on all 14 local and 14 full-field cluster means. If not stated otherwise, the non-parametric Wilcoxon signed-rank test was used for statistical testing (Fig. 3-5, SFig. 2-5).

<u>Pharmacology</u> – To analyse drug-induced effects on BC clusters (Fig. 4, SFig. 4), response traces and RFs of ROIs in one recording field belonging to the same cluster were averaged if there were at least 5 ROIs assigned to this cluster. Spatial RFs were aligned relative to the pixel with the highest s.d. before averaging.

<u>Centre-surround properties</u> – Ring maps of individual ROIs were aligned relative to its peak centre activation and averaged across ROIs assigned to one cluster. For isolating BC surround, centre rings (first 2 rings) were cut and the surround time and space components were extracted by singular value decomposition (SVD). The surround space component was then extrapolated across the centre by fitting a Gaussian and an extrapolated surround map was generated. To isolate the BC centre, the estimated surround map was subtracted from the average map and centre time and space components were extracted by SVD. Estimated centre and surround maps were summed to obtain a complete description of the centre-surround structure of BC RFs. Across clusters, the estimated centre-surround maps captured 93.7 ± 1.4% of the variance of the original map.

The 1-dimensional gauss fits of centre and surround space activation were used to calculate centre and surround ratios (CSRs) for various stimulus sizes. Specifically, the CSR was defined as

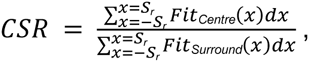

where *S*_*r*_ corresponds to the stimulus radius and ranged from 10 to 500 μm. Time kernels for different stimulus sizes were generated by linearly mixing centre and surround time kernels, weighted by the respective CSR.

<u>BC spectra</u> – Temporal spectra of BC clusters were calculated by Fourier transform of the time kernels estimated for a local (100 μm in diameter) and full-field (500 μm in diameter) light stimulus (see Centre-surround properties). Due to a lower S/N ratio of time kernels estimated for the full-field stimulus, kernels were cut 100 ms before and at the time point of response, respectively, still capturing 86.7 ± 14.7% of the variance of the original kernel. The centre of mass (*Centroid*) was used to characterise spectra of different stimulus sizes and was determined as

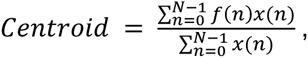

where *x(n)* corresponds to the magnitude and *f(n)* represents the centre frequency of the *n^th^* bin.

<u>Surround chirp and spot noise data</u> – To investigate the effect of surround-only activation and stimulus size on temporal encoding properties across BC clusters, response traces and estimated kernels of ROIs in one recording field belonging to the same cluster were averaged if there were at least 5 ROIs assigned to this cluster. Spectra for kernels estimated from local and full-field spot noise stimuli were calculated as described above.

<u>Time kernel correlation</u> – To analyse the similarity of temporal kernels estimated for a specific stimulus size (Fig. 5l,m), we computed the linear correlation coefficient of each kernel pair from clusters of the same response polarity. We then calculated the average correlation coefficient for every cluster (Fig. 5l) and further across all cluster averages (Fig. 5m).

## ACKNOWLEDGMENTS

We thank G. Eske for technical support, C. Behrens for help with EM data, J. Jüttner from the B. Roska lab (FMI for Biomedical Research, Basel) for help with the virus injection protocol and X. Pitkow and R. Taylor for feedback on the manuscript. We thank Loren L. Looger, the Janelia Research Campus of the Howard Hughes Medical Institute and the Genetically-Encoded Neuronal Indicator and Effector (GENIE) Project for making the viral constructs (AAV9.hSyn.iGluSnFr.WPRE.SV40, AAV9.CAG.Flex.iGluSnFr.WPRE.SV40 and AAV9.Syn.Flex.GCaMP6f.WPRE.SV40) publically available. This work was supported by the Deutsche Forschungsgemeinschaft (DFG; Werner Reichardt Centre for Integrative Neuroscience Tübingen, EXC307 to M.B., T.S. and T.E; BA 5283/1-1 to T.B; BE 5601/1-1 to P.B.), the German Federal Ministry of Education and Research (BMBF; BCCN Tübingen FKZ 01GQ1002 to M.B and T.E), the BW-Stiftung (AZ 1.16101.09 to T.B.), the intramural *f*ortμne program of the University of Tübingen (2125-0-0 to T.B.), and the National Institute of Neurological Disorders and Stroke (U01NS090562 to T.E.) as well as the National Eye Institute (1R01EY023766 to T.E.) of the National Institutes of Health. The content is solely the responsibility of the authors and does not necessarily represent the official views of the funders.

## AUTHOR CONTRIBUTIONS

KF, PB, TE and TB designed the study; KF performed experiments and pre-processing; KF
performed viral injections with help from TS; PB developed the clustering framework with
input from MB; KF, PB and TB analysed the data with input from TE; KF, PB, TE and TB
wrote the manuscript.

## SUPPLEMENTARY DATA LEGENDS

**SVideo 1 ∣ Glutamate responses in the IPL**. Background-subtracted and colour-coded glutamate signals recorded in the On ChAT band (64×16 pixels@31.25 Hz), where yellow corresponds to higher activity. Responses to the noise stimulus (exemplary 30 s, single trial) and local chirp (mean across n=5 trials) are shown. Scan field corresponds to the one shown in (Fig. 1d). Video in real-time.

**SFigure 1.**
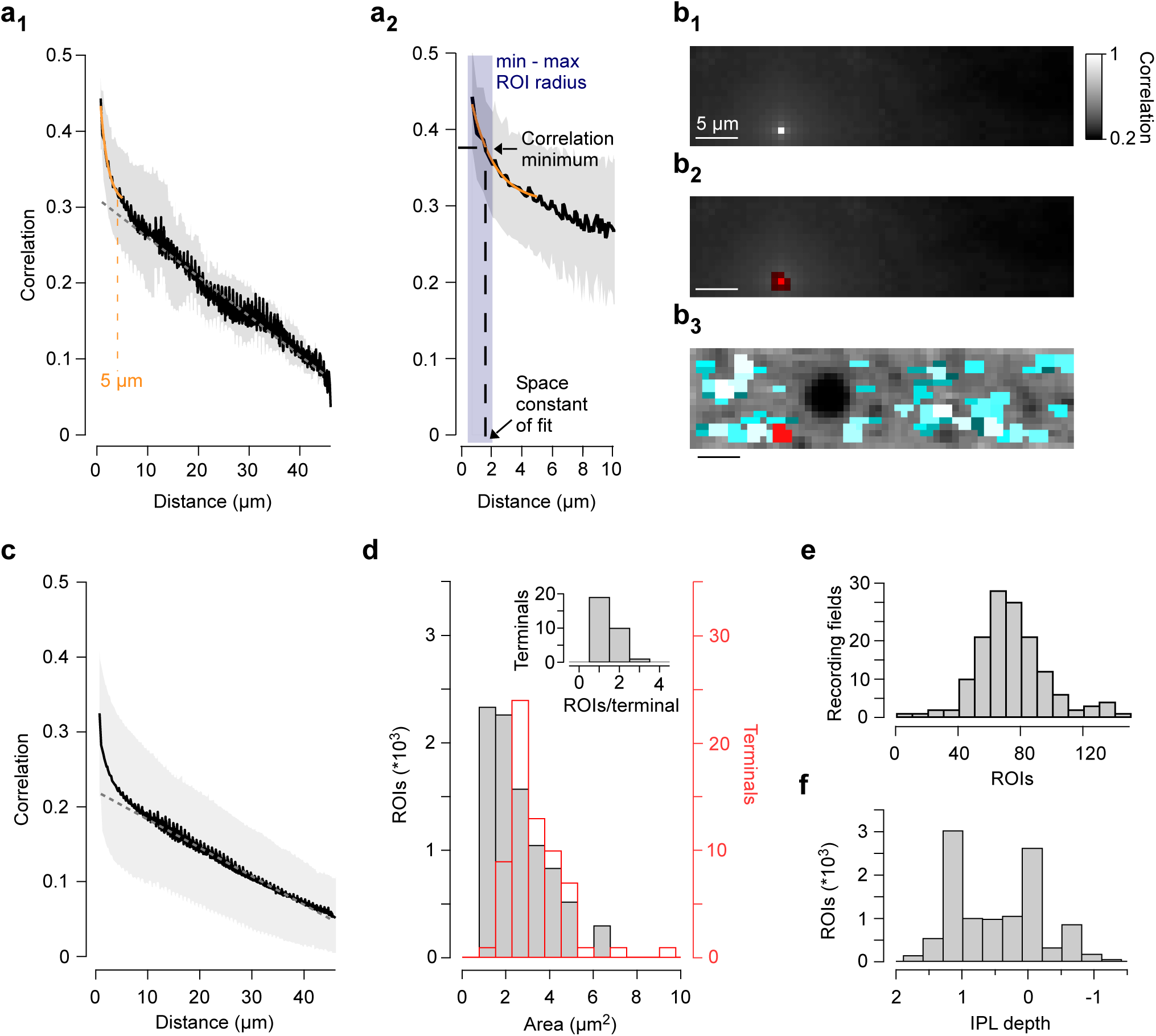
ROI detection. **a**, Mean correlation (± s.d. shading, n=100 pixels) between noise-response traces of two individual pixels from scan field shown in (Fig. 1d) plotted against the distance of each pixel-pair (a_1_). Dotted line shows linear fit to the data above x = 10 μm and its extrapolation towards x = 0 μm. The space constant obtained from an exponential fit (*orange*) for distances >5 μm was used to determine a scan field’s specific correlation minimum for ROI detection (a_2_, zoomed-in version of a_1_). Blue shading indicates the range of allowed ROI radii (0.375-2 μm). **b**, Scan field from (Fig. 1d), with each pixel color-coded according to its correlation with the noise trace of pixel indicated in (b_1_). In b_2_, the red shading corresponds to pixels with a correlation coefficient > correlation minimum from (a_2_), resulting in red ROI in (b_3_). **c**, As (a_1_), averaged for n=71 scan fields recorded at 48x12 μm. **d**, Histogram of ROI (*black*) and BC axon terminal (*red*) area. Terminal area was determined from BC axonal arborisations labelled in GCaMP6-injected PCP2 mice where individual axon terminals can be distinguished (cf. SFig. 2a). Inset shows a histogram of the number of ROIs per BC axon terminal. **e**, Distribution of ROI numbers per scan field. **f**, Histogram illustrating sampling of ROIs against IPL depth. Data collected specifically for drug experiments (cf. Fig 4; Methods) contributed to the “oversampling” of the two ChAT bands.

**SFigure 2.**
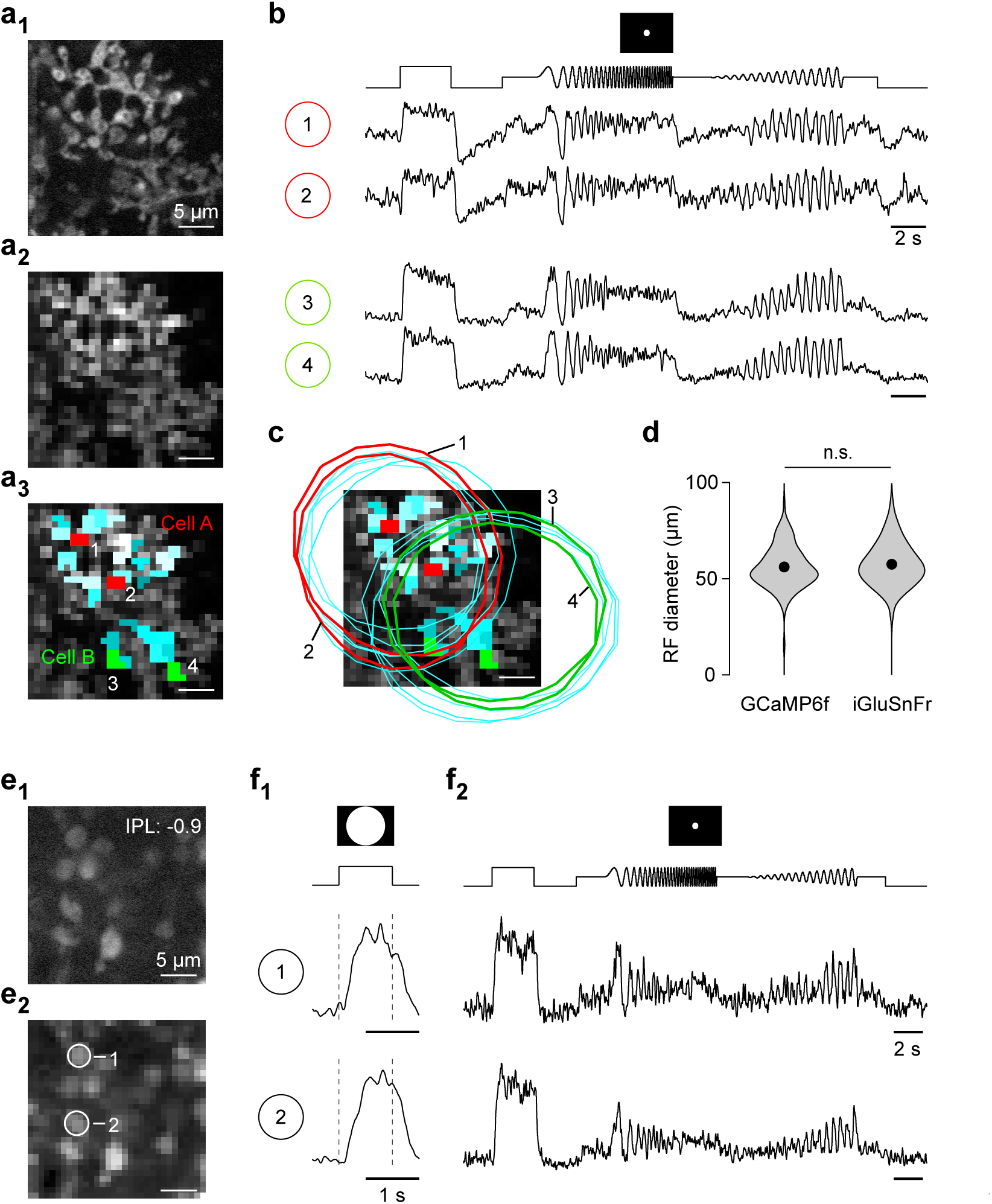
GCaMP6 signals in mouse BC axon terminals. **a**, High resolution scan of GCaMP6f-expressing BC axon terminal systems in the IPL of a Pcp2 mouse (a_1_) and corresponding scan field (a_2_) with automatically generated ROI mask overlaid (a_3_, cf. SFig. 1). **b**, Exemplary mean local-chirp responses (n=5 trials) of individual ROIs shown in (a_3_) of the two different BCs shown in a_1_. **c**, Scan field from (a_3_) with ROI mask and spatial RFs (2 s.d. outlines of gauss fit) overlaid. **d**, Distribution of RF diameters estimated from On BC terminal calcium (GCaMP6) and glutamate release (iGluSnFr), respectively. Black dots correspond to mean RF diameters (56.1 ± 10 μm for GCaMP6 and 57.5 ± 10.6 μm for iGluSnFr). p>0.05, n=261 (GCaMP6) and n=3,540, non-parametric non-paired Wilcoxon signed-rank test. **e**, High resolution scan of GCaMP6-expressing RBC axon terminals (e_1_) and corresponding scan field (e_2_), with two individual terminals indicated. **f**, Mean responses of terminals shown in (e_2_) to full-field flashes (f_1_, n=20 trials) and local chirp stimulus (f_2_, n=5 trials).

**SFigure 3.**
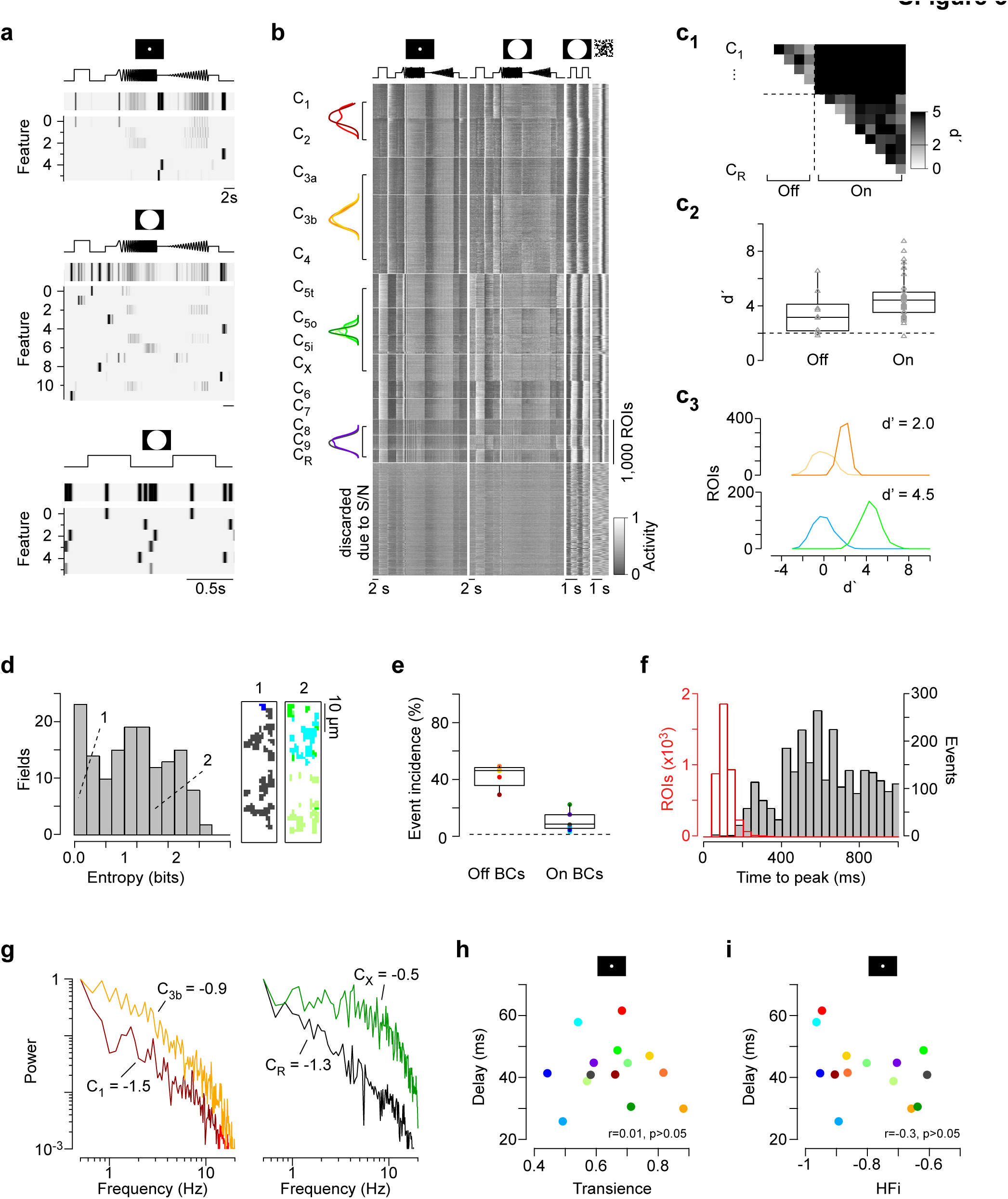
Clustering. **a**, Temporal features extracted from glutamate traces in response to local (n=6 features, *top*) and full-field (n=12 features, *middle*) chirp and full-field flashes (n=6 features, *bottom*). **b**, Heat maps of all recorded glutamate responses (n=14 clusters plus ROIs discarded based on signal-to-noise (S/N) ratio) to the four visual stimuli (cf. Fig. 1); n=11,101 ROIs from 29 retinas. Each line corresponds to the responses of individual ROIs with activity colour-coded. Block height represents the number of included ROIs per cluster. Within one cluster, ROIs are sorted based on the quality of their RFs and local chirp response (Methods). Overlaid stratification profiles (*left*) illustrate overlap for some types. **c**, Cluster separation was determined for every cluster pair using the sensitivity index d’ (c_1,2_). Dotted lines in (c_1_) illustrate transition between On and Off clusters and dotted line in c_2_ at d’=2, which corresponds to about 15% FP/FN rates. c_3_, Separation of exemplary cluster pairs with a low (d’=2.0, *top*) and an average (d’=4.5, *bottom*) sensitivity index, respectively. **d**,. Distribution of field entropies (*left*, Methods). Two exemplary scan fields (*right*) with ROIs colour-coded by cluster allocation illustrate low (*1*) and high (*2*) field entropy, respectively. **e**, Percentage of ROIs with at least one opposite polarity event in response to the local chirp step response for On and Off BC cluster. Dotted line illustrates mean incidence of spontaneous events (<1%). **f**, Time to peak of On events observed in the Off layer (*grey*) and On responses in On layer (*red*). Events were estimated from responses to single trials of the local chirp stimulus. **g**, Mean spectra (n=5 trials) of local chirp step responses for two Off (*left*) and two On (*right*) ROIs shown in (Fig. 2e), with HFi estimated from the relative power of low (0.5-1 Hz) and high (2-16 Hz) frequencies (Methods). **h**, Mean response transience of BC clusters is not correlated with mean response delay. r=0.01, p>0.05, n=14, linear correlation. **i**, Mean HFi and mean response delay are not correlated across BC clusters. r=0.3, p>0.05, n=14, linear correlation.

**SFigure 4.**
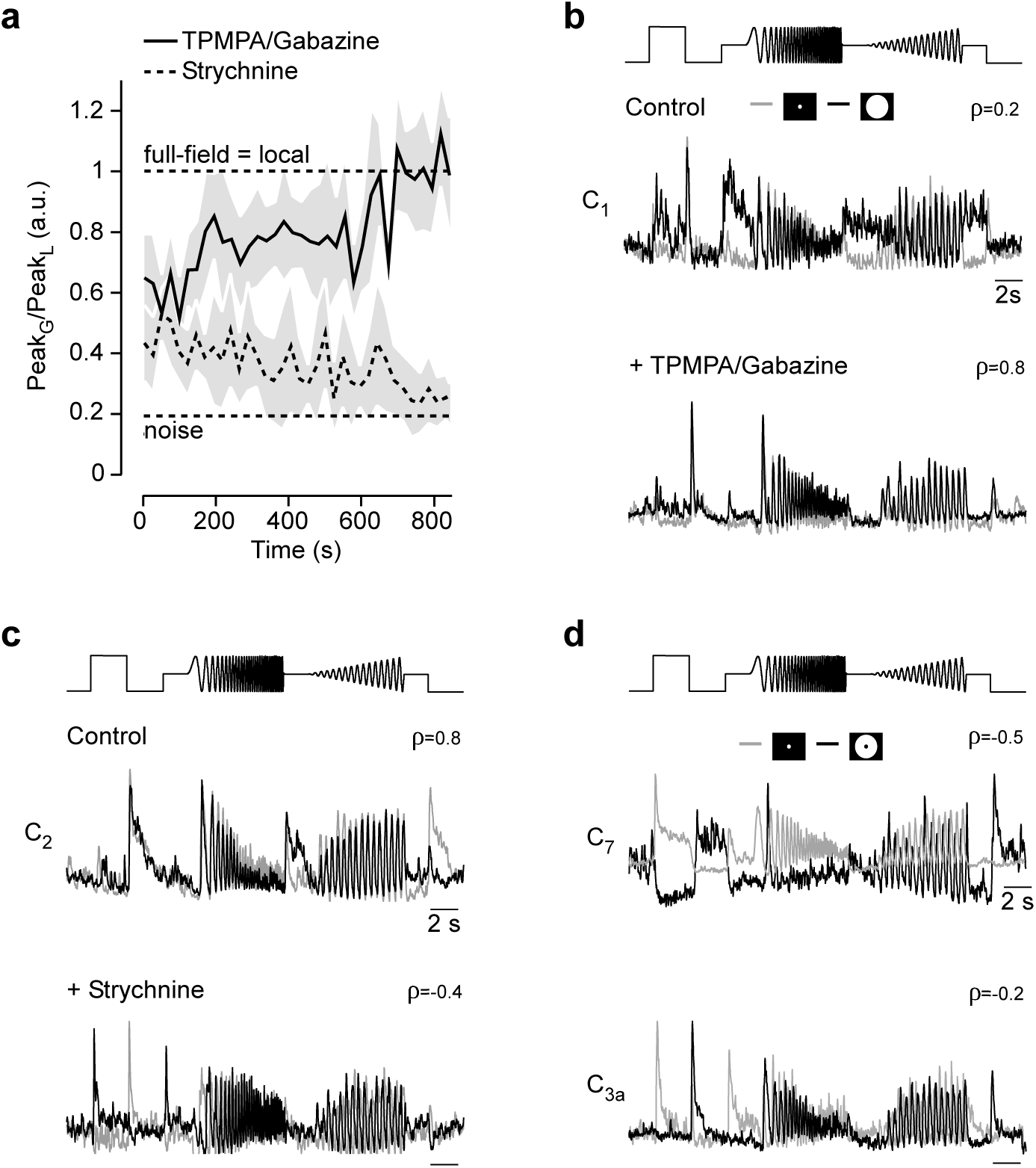
GABA and glycinergic inhibition differentially shapes BC responses. **a**, Change of mean peak amplitudes (± s.d. shading) of n=15 ROIs originating from two scan fields during wash-in of GABA and glycine receptor blockers, respectively (cf. Fig. 4c). **b,c**, Local (*grey*) and full-field (*black*) chirp responses for control and drug conditions (b: GABA receptor block; c: glycine receptor block), with linear correlation coefficient (ρ) between each pair indicated. **d**, Local (*grey*) and surround (*black*) chirp responses for an exemplary On (C_7_) and Off (C_3a_) BC cluster, respectively.

**SFigure 5.**
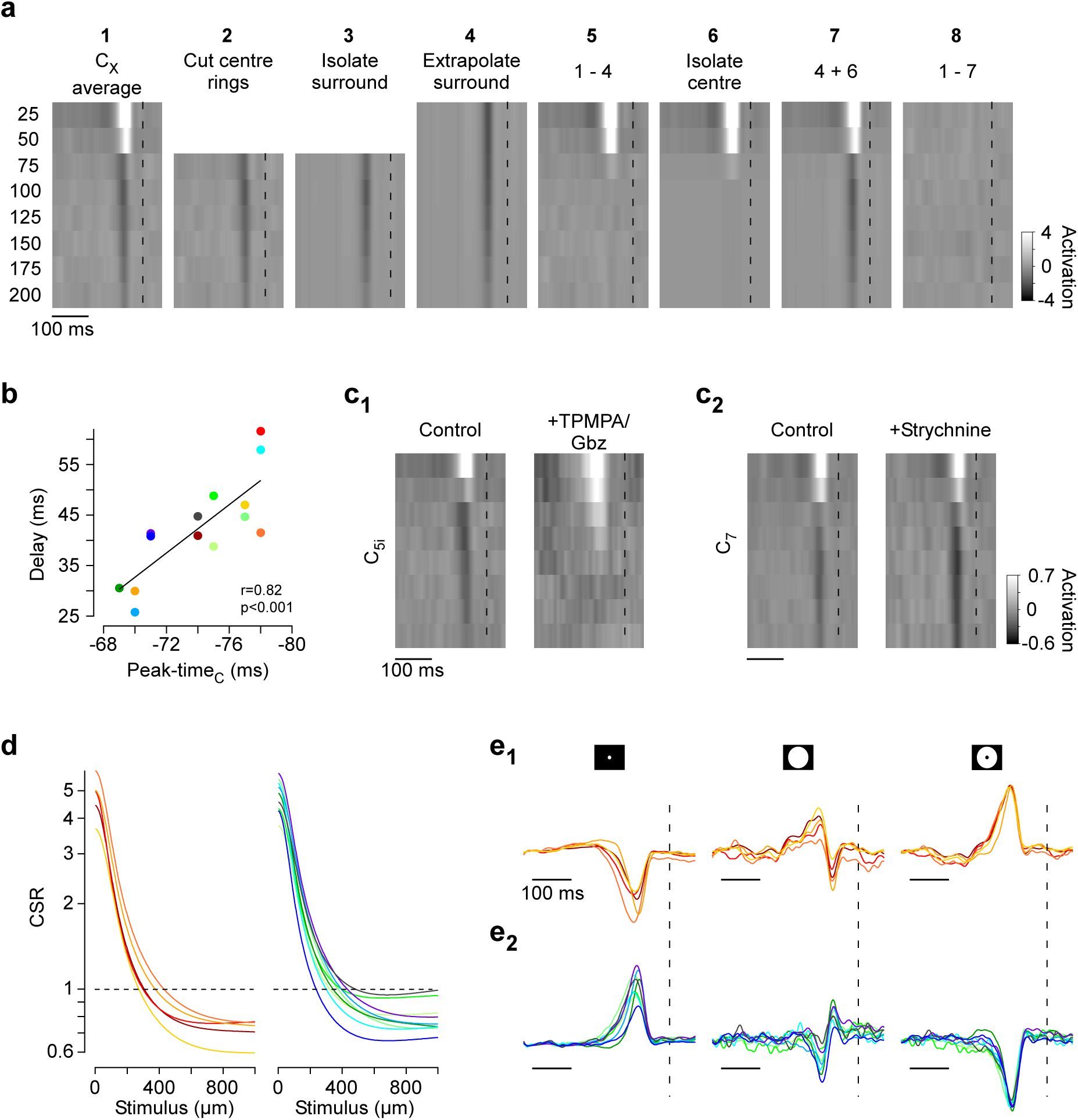
Centre-surround RFs of BC clusters. **a**, Centre-surround maps obtained from the ring noise were averaged across ROIs assigned to one cluster (*1*). To isolate the surround component, the innermost two rings which contained the majority of the centre component were clipped (*2*) and the surround map estimated by SVD (*3*) was extrapolated across the centre by fitting a Gaussian (*4*). Next, the extrapolated surround map was subtracted from the average map and the centre component was extracted using SVD (*6*). The resultant centre-surround map was then subtracted from the average map to estimate the residual variance (*7*, Methods). Dotted lines at t=0. **b**, Time to peak of centre time kernels correlated with response delay estimated from local chirp step responses (r=0.82, p<0.001, n=14, linear correlation), indicating that centre kernels adequately reflect BC responses to local stimuli **c**, Effect of GABA (c_1_) and glycine (c_2_) receptor block on centre-surround RFs of two exemplary BC clusters. Centre-surround maps correspond to averages on n>5 ROIs of one scan field. **d**, Predicted centre-surround ratios (CSRs) of BC clusters for different stimulus diameters (cf. Fig. 5d). **e**, Normalised temporal kernels predicted for local (100 μm diameter), full-field (500 μm diameter) and surround-only (500 and 100 μm outer and inner diameter, respectively) stimulation for Off (*top*) and On (*bottom*) BC clusters (cf. Fig. 5e), respectively.

